# Observation of electromyogram and electrocardiogram changes with deqi for the development of an objective evaluation index of the occurrence of deqi

**DOI:** 10.1101/2023.10.18.562998

**Authors:** Nao Miyamoto, Yusuke Sakaue, Shima Okada, Naruhiro Shiozawa

## Abstract

A unique sensation called “deqi” occurs in acupuncture, which is considered an essential variable in the study of the mechanisms and effects of acupuncture. However, an objective method for evaluating deqi is yet to be established. Therefore, we aimed to create an objective evaluation index for deqi using electromyogram (EMG) and electrocardiogram (ECG), which are expected to present characteristic changes based on the findings of previous studies. This study included 16 healthy adult male subjects. ST36 stimulation was performed three times after a one-week washout (N=48). EMG was used to calculate myopotentials, and ECG was used to calculate the heart rate (HR) immediately after stimulation. The subjects declared whether Deqi occurred, and the characteristic reactions of deqi in EMG and ECG were evaluated. In 48 trials, Deqi occurred 42 times and did not occur 6 times. Myopotentials were significantly higher when deqi occurred than when it did not. Furthermore, a positive correlation was identified between subjective sensation and the rate of change in myopotential. Deqi significantly increased the myopotential levels, indicating that it induced involuntary muscle activity. This involuntary muscle activity may be related to the stretch reflexes and nerve stimulation. For the HR immediately after stimulation, no change with deqi was identified, although a significant decrease in HR was found in cases without deqi. Deqi could induce a transient stress response and inhibit the decrease in the HR caused by acupuncture. The results of this study indicate two characteristics of deqi obtained from EMG and ECG: an increase in myopotential and an inhibition of the decrease in HR expected with acupuncture. These characteristics could provide objective evidence for the occurrence of deqi. Future studies with larger sample sizes are required toto further strengthen the objective assessment of deqi.

## Introduction

Acupuncture is a medical intervention that employs extremely fine needles to stimulate specific points on the body surface, i.e., acupoints, that are believed to possess therapeutic efficacy. Deqi, a complex sensation that is primarily felt beneath the skin, occurs during acupuncture and is considered vital in traditional Chinese medicine because it is a requirement for therapeutic efficacy and a prognosis predictor [1]. However, the reason for which deqi is essential and its underlying mechanisms remain unelucidated [2]. Although the definition of deqi varies among studies, it is primarily described as the subjective feelings and perceived reactions of both the patient and the acupuncturist [2–6]. However, no clear standards exist for distinguishing the occurrence of deqi. Previous studies have assessed the sensations experienced by patients [7]; one method for such assessment is the Acupuncture Sensation Questionnaire, which is mainly used to classify deqi sensations [5]. However, this questionnaire does not include distinct criteria for determining the occurrence of deqi; therefore, limitations exist when using solely subjective evaluations. Hence, the first step to elucidate the mechanisms of deqi is to find an objective method for detecting it.

One of the characteristic biological responses to deqi is an increase in myopotential at the stimulus’ localization; significant increases in myopotential as deqi occurs have been reported [8–10]. This suggests that deqi affects peripheral motor nerves. Another characteristic change was observed in heart rate (HR); acupuncture stimulation usually induces somatic autonomic reflexes, which reduce HR by inhibiting cardiac sympathetic nerve activity [11,12]. However, the HR does not decrease immediately after stimulation when deqi occurs [13]; this suggests that a stress response occurs immediately after stimulation when deqi takes place, which may suppress the decrease in HR [14]. The HR increase in the stress response to an invasive stimulus occurs within a very short period immediately after it [15,16]. Previous studies have evaluated the effects of acupuncture stimulation on HR by calculating averages of 2–5 min, whereas transient responses to deqi were observed in a much shorter period.

Accordingly, we formulated the following two hypotheses: First, a significant increase in myopotential occurs only when deqi occurs. Second, the HR increases temporarily in response to stress immediately after deqi occurs. We aimed to determine whether the occurrence of deqi could be objectively assessed using electromyogram (EMG) and electrocardiogram (ECG). Therefore, we stimulated Zusanli (ST36), which is the most widely employed acupoint in research, and compared the myopotentials obtained from EMG and the change in HR obtained from ECG with and without the presence of deqi, based on the subject’s declaration of such. This study provides evidence that it is possible to objectively assess the occurrence of deqi by using EMG and ECG.

## Methods

### Participants

This study included 16 healthy adult males. Females were excluded because their menstrual cycle affects HR. Subjects with skin diseases were excluded since acupuncture needles were inserted through the skin. We also excluded subjects with cardiac disease, as it would affect the measurement of HR. In addition, the subjects selected have had previous experience with acupuncture to prevent fear or anxiety from influencing HR. The purpose of the study, freedom to participate or discontinue the study, protection of personal information and measurement data, and possible adverse events were clearly explained to the participants, and their informed consent with written was obtained. Subject recruitment began in March 2021 and ended in September 2021. This study was approved by the Ethics Review Committee of the Ritsumeikan University (February 22, 2020, Approval No. BKC-人医-2020-059).

### Experimental protocol

First, an ultrasound examination was performed to determine the anatomical location of ST36, which was then marked. Next, preparations for ECG and EMG measurements were performed as follows: the subject was then placed supine on the treatment table for 20 minutes to stabilize his condition; measurements were performed 5 minutes before stimulation. After measurement, the needle was inserted 5 mm and held in that position for 1 minute, to separate deqi from pain. The needle was then inserted at a rate of 1–2 mm per second, and the subject was instructed to press a switch when he felt deqi; the switch was used to synchronize deqi with the ECG or EMG signals. If deqi occurred, the needle was held at the depth declared by the subject, and if no declaration was made, the needle was inserted up to 30 mm; the maximum depth. After confirming the subject’s declaration of deqi, the needle was maintained in place for 5 minutes post-measurement. Finally, the experiment was terminated when the participant responded to the Massachusetts General Hospital Acupuncture Sensation Scale (MASS, Fig 1).

**Fig 1.**
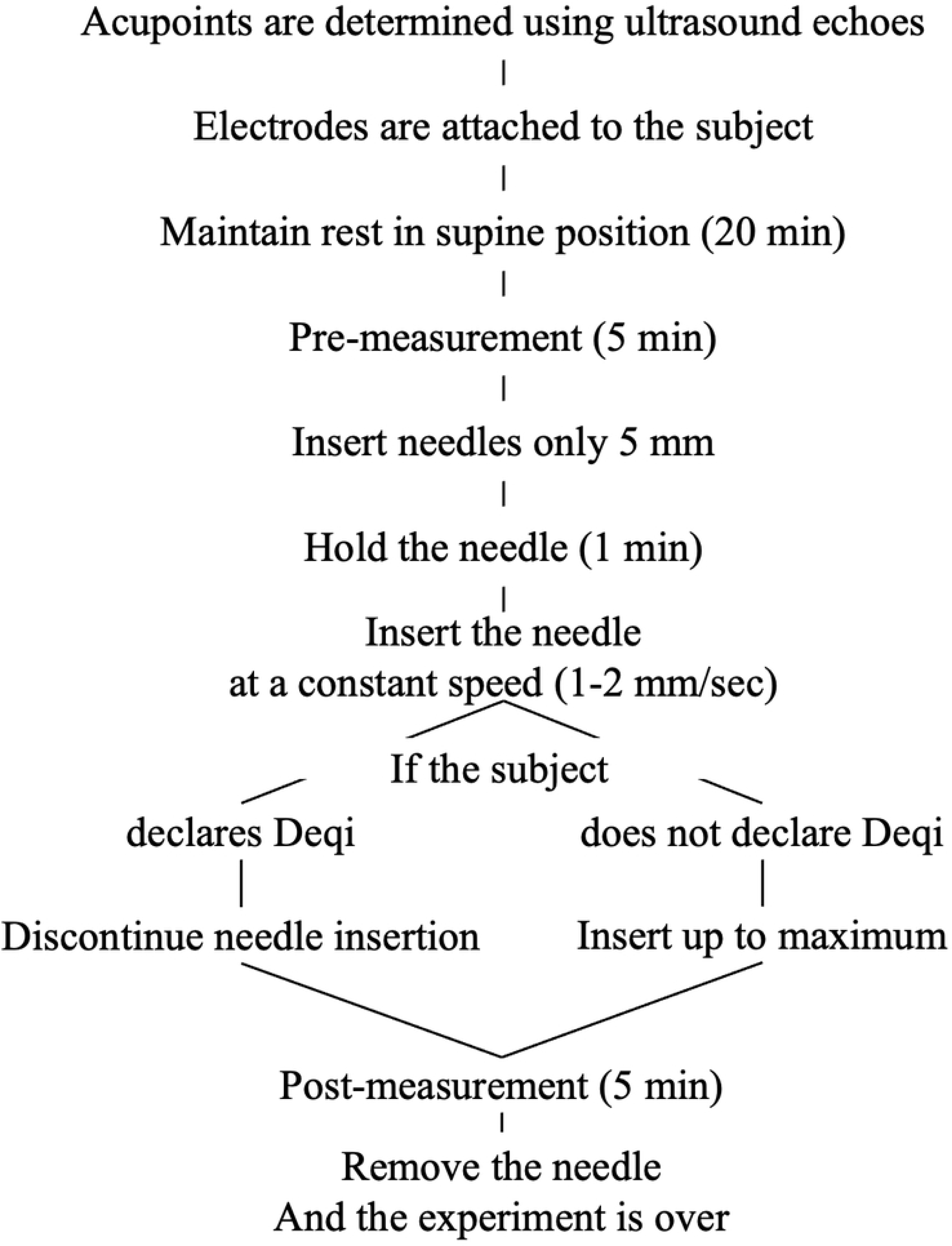
Experimental Protocol.

The above described protocol was performed three times on the same day of the week, with a six-day washout period in between, as the effect of acupuncture was considered to disappear after six days in previous studies [17]. To account for diurnal variation, the subjects started measurements at the same time of the day on all three days. We divided the two groups to examine the usefulness of EMG and HR as objective indicators of deqi, as follows: days with deqi were designated as the acupuncture with deqi group (AWD); and days without deqi were designated as the acupuncture without deqi group (AOD).

### Subjective evaluation

We used the Japanese version of the MASS, developed by the Massachusetts General Hospital, to investigate acupuncture sensation; such was created by Nishiwaki et al. in 2017 and has been reviewed for authenticity. The MASS consists of 13 items (including soreness, aching, deep pressure, heaviness, fullness/distension, tingling, numbness, sharp pain, dull pain, warmth, cold, and throbbing)[18]. The questions can be answered on an 11-point scale (0 = none, 10 = unbearable) to evaluate acupuncture sensation, even if deqi does not occur. The total score for each question was used to determine the strength of deqi.

### Objective evaluation

The EMG and ECG signals were measured using a Biosignal Amplifier System (Polymate V AP5148, Miyuki Giken, Tokyo, Japan). All the electrodes used in the measurements were small active electrodes (AP-C151; Miyuki Giken, Tokyo, Japan).

EMG was measured using the anode of the electrode fixed 30 mm above ST36, the cathode affixed 30 mm below ST36, and the ground affixed to the outer margin of the ankle joint (fibula). The sampling frequency was set to 4 kHz. After the electrodes were applied, the subjects were asked to flex and extend their ankle joints to check whether the EMG was measured normally. Myopotentials were calculated by processing the root mean square of the electrical signals obtained from the EMG within a specific time range. Noise was defined as the median 0.5 sec of the values measured at rest before stimulation. The signal was also defined in two patterns, depending on the presence or absence of deqi: 1) When deqi occurred, the analysis range was 0.25 s before and 0.25 s after the declaration of deqi (0.5 sec in total)―as the human reaction time is limited, this range also includes the time before the declaration; 2) When deqi did not occur: 0.5 s after reaching the maximum depth was defined as the signal.

The signal-noise ratio (SNR) was calculated by dividing the obtained signal by the noise [12]. Deqi-induced muscle activity was expected to be microscopic; hence, SNR was used to assess muscle activity. A change in the SNR > 1 indicated that some motor nerves were activated, as electrical activity occurred more frequently than at rest.

For the ECG, the cathode was placed on the left sixth rib of each subject, anode on the right sixth rib, and ground electrode on the outer left sixth rib. The signals were low-pass filtered at 60 Hz before amplification and recorded at a sampling frequency of 1 kHz. The R-R interval was calculated using the R waveform obtained from the ECG. HR-pre was defined as the average of the measurement 5 min before stimulation, whereas HR-post was divided into two patterns depending on the presence or absence of deqi: 1) With deqi onset, 90 s after the declaration was used as the analysis interval; 2) In the absence of deqi, the maximum analysis interval was 90 s after insertion of the needle to the maximum depth.

In both cases, the average R-R interval was calculated every 30 s and converted to the HR; this was designated as the HR-post. The data obtained were normalized by dividing the HR-post by the HR-pre, and this was defined as the rate of change in HR. An HR change rate > 100 indicated an increase in HR.

### Acupuncture stimulation

The Acupoint stimulated was Zusanli (ST36) on the right side, which was located below the knee and above the anterior tibialis muscle. The location of ST36 was defined as three cun (cun is a unit of length used in acupuncture that is unique to traditional Chinese medicine) below the calf on a line connecting ST35 and ST41 on the anterior surface of the lower leg, with the deep fibular nerve and anterior tibial artery below (Fig 2A) [19]. In this study, ST36 was defined as being directly above the anterior tibial artery. The ST35 and ST41 levels were measured on the body surface; we first marked these two points with a pen and then measured the distance between the two points, divided it by 16 and multiplied the result by 3 to obtain the height of ST36. At the defined height, ultrasonography was performed using a ProSound SSD-3500 (Hitachi Aloka Medical, Tokyo, Japan) to determine the anatomical location of ST36. The probe (L3-12-D, GE Healthcare Japan) was applied to the target site with the upper surface of the echo image facing the knee joint and the lower surface facing the ankle joint. Measurements were made using ultrasound B-mode (Fig 2B).

**Fig 2.**
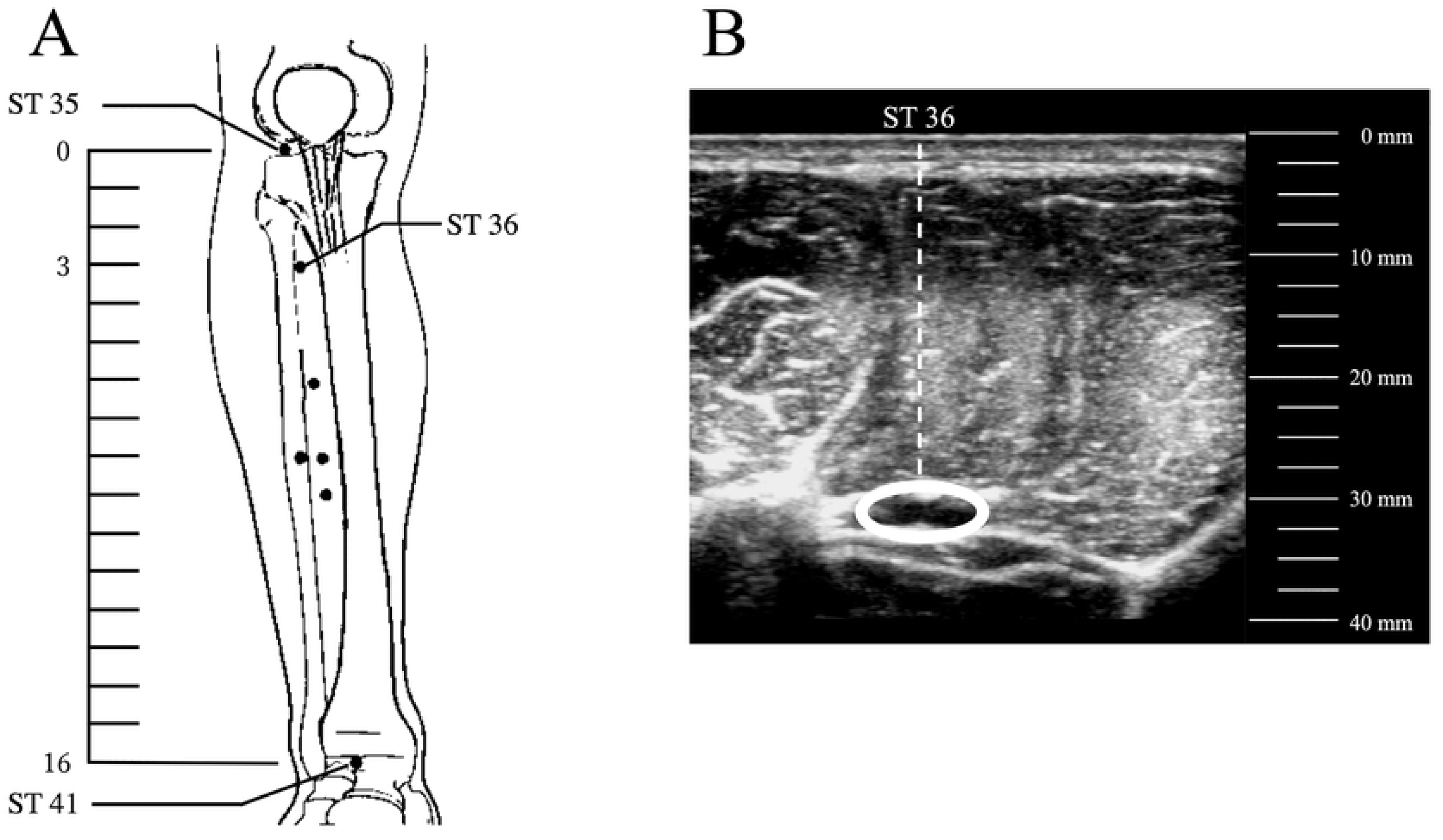
Zusanli (ST36) being located and the corresponding ultrasound image. (A) Definition of the location of ST36 is shown. (B) Ultrasound imaging of ST36 shows that the artery is present.

Disposable stainless steel needles (length, 40 mm; thickness,: 0.20 mm; Seirin, Shizuoka, Japan) were used for stimulation. The angle of the needle was maintained perpendicular to the skin surface, and the needle was inserted monotonically at a constant speed (1–2 mm/sec) without any unique manipulation. The needle was inserted to the maximum depth at which the anterior tibial artery was located.

### Statistical analysis

Student’s t-tests (at a significance level of 5%) were performed between AWD and AOD for noise, signals, and SNR. Two-way repeated-measures analysis of variance was performed to investigate the differences in HR for the two factors of group and time. Dunnett’s test was used as the post-hoc test if a main effect was found over time, whereas Welch’s t-test was used as the post-hoc test if a main effect was found for the group. Statistical analysis was performed on the overall data (n=48) using Spearman’s correlation coefficient to examine whether a relationship between the subjective assessment (strength of Deqi; MASS total score) and the objective data (SNR, HR; only 30 s) was present. Statistical analyses were performed using statistical analysis software R (version 4.1.1, R Foundation for Statistical Computing, Vienna, Austria)[20].

## Results

The 16 subjects were 21.9±1.9 years old, with a height of 170.5±4.1 cm, a weight of 66.5±9.7 kg, and a body mass index 22.9±3.1 kg/m^2^. In this study, no subjects dropped out and no adverse events associated with acupuncture stimulation were identified. Three stimulation sessions were provided to 16 subjects, and data for 48 sessions were obtained.

### Subjective evaluation

Deqi occurrence was induced 42 times in the AWD group (N=42); deqi was not induced 6 times in the AOD group (N=6; Table 1). All participants declared deqi at least twice. The strength of deqi was 23.7 ± 14.2 in AWD and 0.5 ± 1.2 in AOD.

**Table 1.**
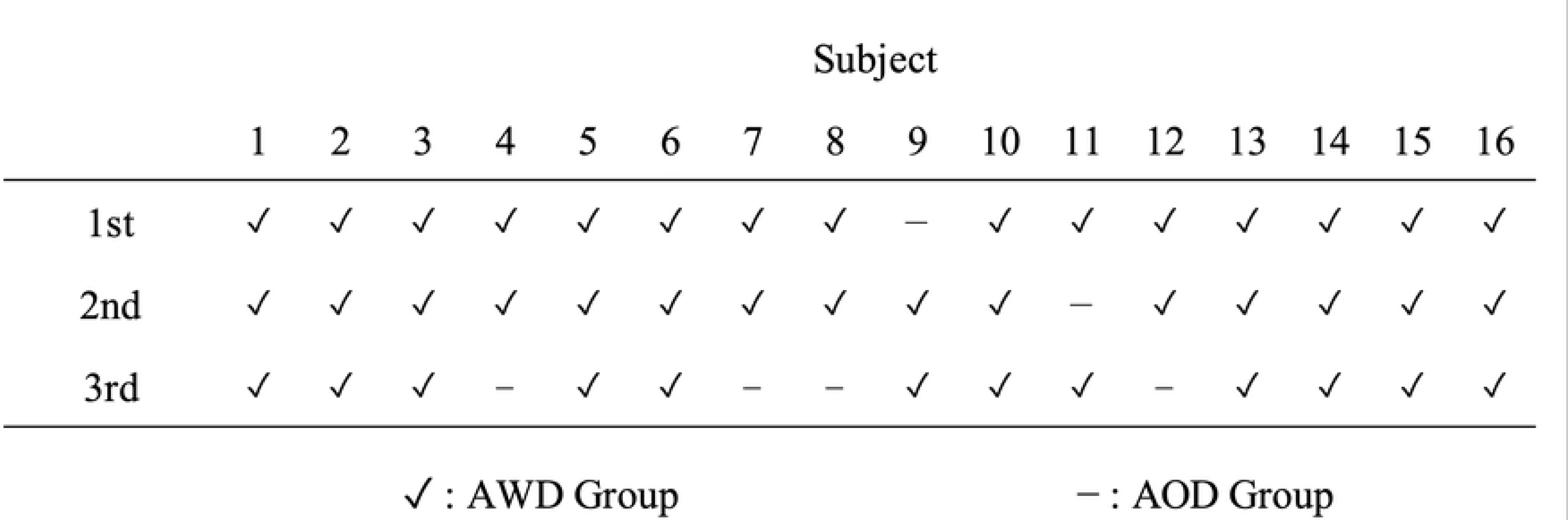
Subject’s Deqi Declaration. 16 subjects were involved the experiment in 3 sessions. The table shows each subject’s declaration of Deqi. Deqi was declared 42 times, and not declared 6 times.

### Objective evaluation

#### Rate of change in myopotential (Figure 3)

**Fig 3.**
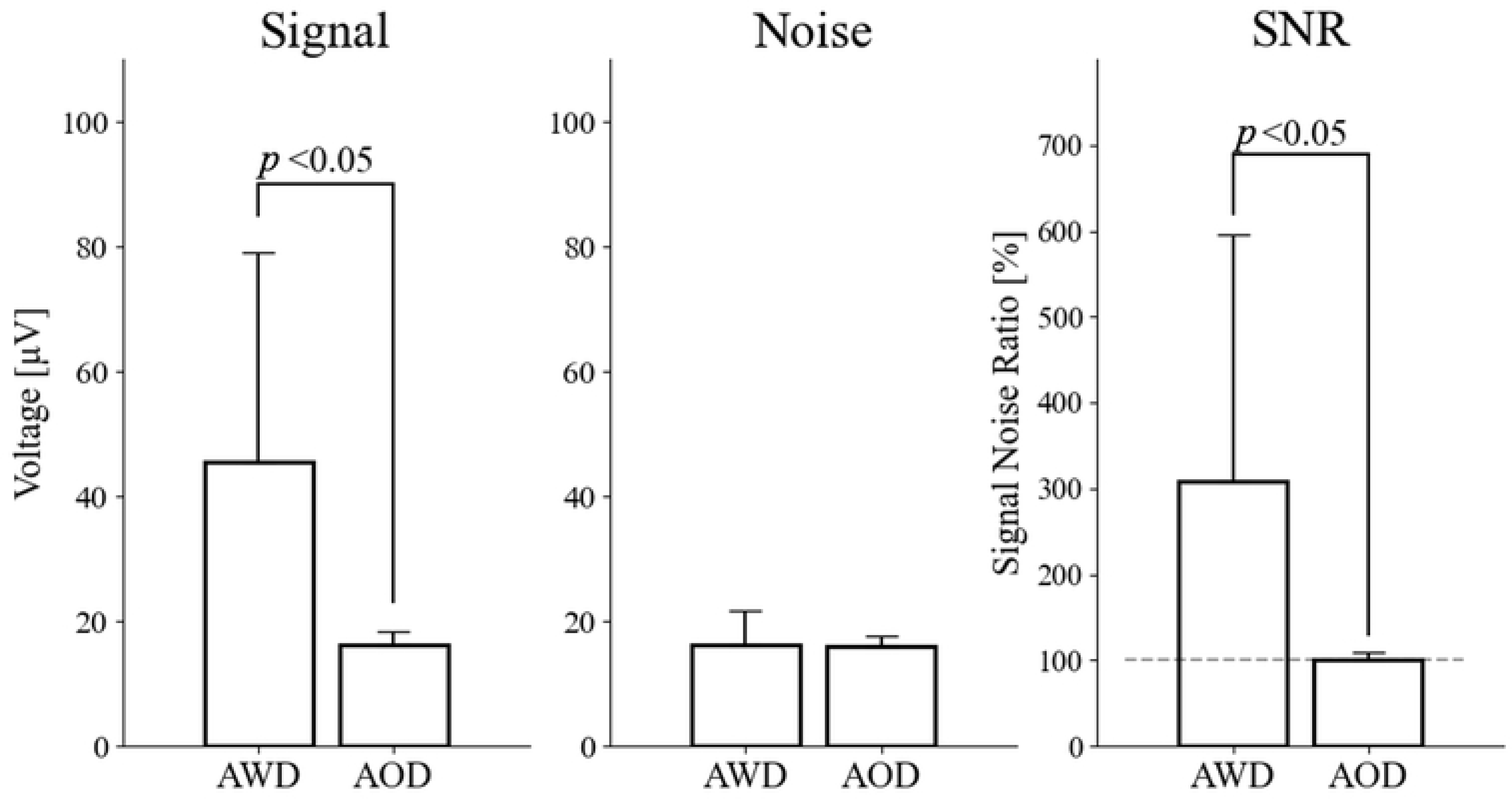
Signal, noise, and signal-noise ratio with and without deqi. Significant differences in the signal to signal-noise ratio are found between presence and absence of deqi. Noise values are not significantly different.

The signal in the AWD was significantly higher than the AOD group (p<0.05). Regarding noise, no significant difference between AWD and AOD was observed. The SNR was significantly higher in the AWD than in the AOD group (p<0.05).

#### Rate of change in HR (Figure 4)

**Fig 4.**
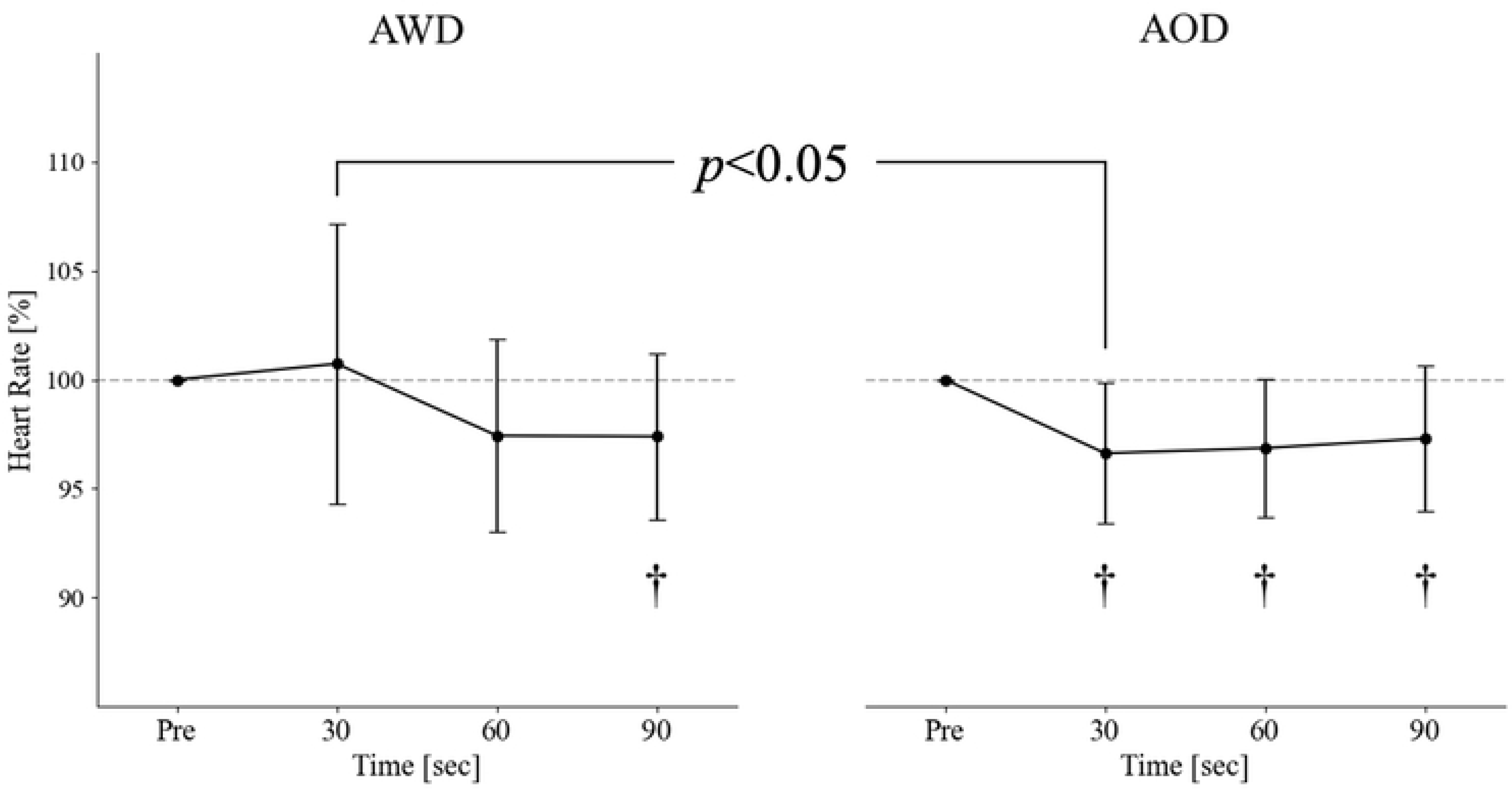
Change of heart rate every 30 seconds in the presence and absence of deqi. With deqi, the heart rate is significantly reduced 90 s after stimulation. Without deqi, the heart rate decreases from the post-stimulus period. Significant differences are found between the two groups 30 s after stimulation.

The results of the repeated two-way analysis of variance showed the main effects of time and group. However, no interaction effects were observed. In the AWD group, a significant decrease in HR was observed relative to HR-pre at 90-s post-stimulation. In the AOD group, a significant decrease in HR was observed relative to HR-pre at 30, 60, and 90-s post-stimulation. The AWD group showed a significantly higher HR at 30 s after stimulation than the AOD.

#### Correlation between subjective and objective evaluation (Figure 5)

**Fig 5.**
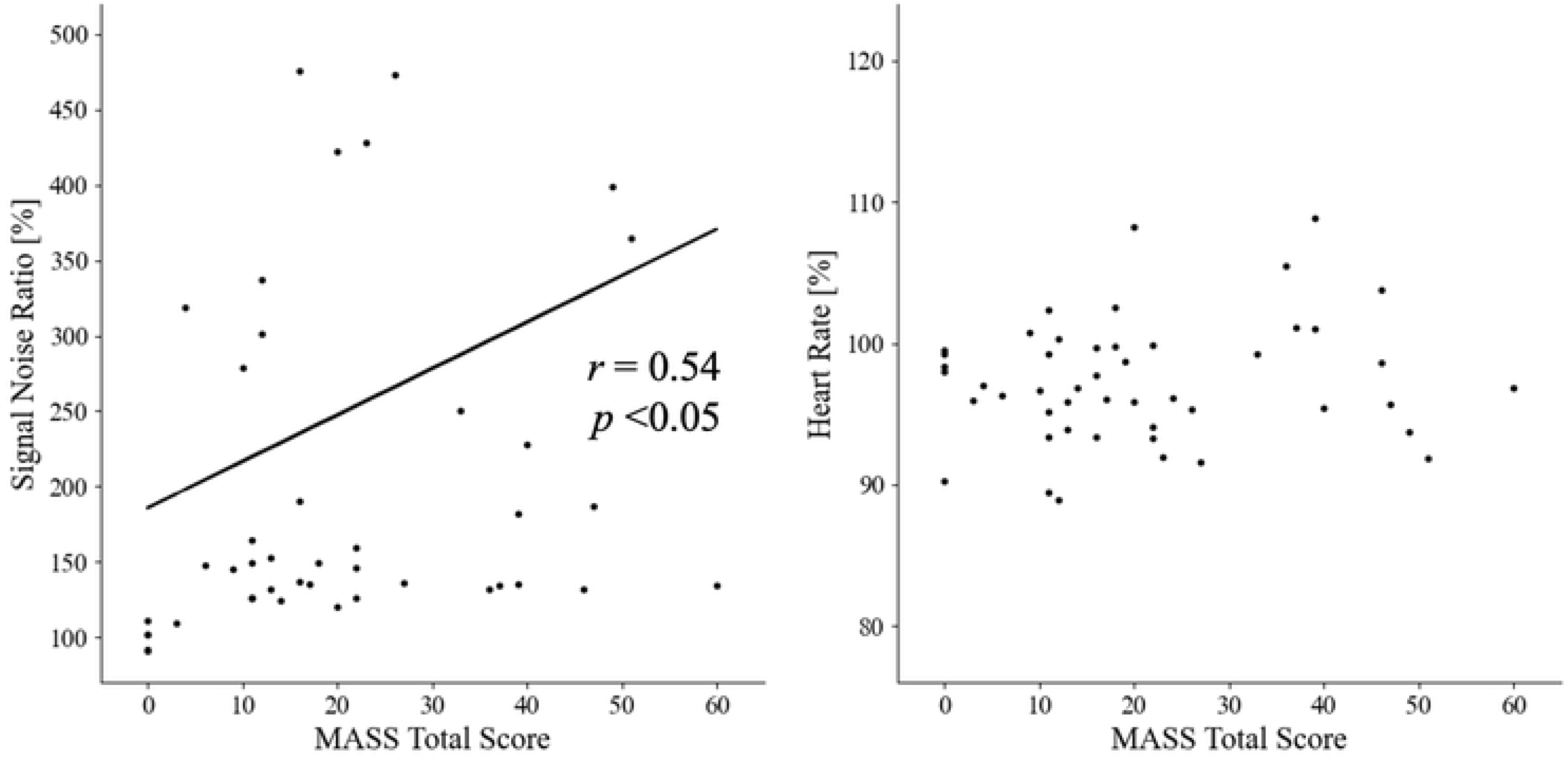
The left figure shows the correlation between myopotentials, and the right figure shows the correlation between heart rate and MASS total points. A significant correlation IS found between EMG potentials and MASS (r=0.54, p<0.05); however, no significant correlation is identified for the heart rate. Abbreviations: MASS, Massachusetts General Hospital Acupuncture Sensation Scale.

A significant positive correlation was found between the subjective evaluation index (strength of deqi) and the SNR (r=0.54, p<0.05). No significant correlation was found between the subjective evaluation index and the rate of change in HR 30 s after stimulation.

## Discussion

The SNR was significantly higher in the AWD than in the AOD group. Furthermore, the average SNR in the AWD group was 309, which was > 100, which indicates that micro muscular activity was induced by deqi. These results provide strong supportive evidence that our first hypothesis was correct. Furthermore, the stronger the perceived acupuncture sensation, the higher the rate of change in myopotential. Several reports have demonstrated an increase in myopotential when deqi occurs, which is similar to the findings of this study [8–10]. In clinical practice, minute muscle contractions are frequently observed when deqi is induced. The muscle contractions triggered by deqi are involuntary reflexes. Consequently, the stretch reflex is considered one of the mechanisms contributing to this involuntary muscle contraction. Muscle spindles, in which intrafusal muscle fibers are arranged parallel to the muscle fibers, are responsible for sensing changes in muscle length and velocity. When a muscle is elongated, the muscle spindle is stretched, increasing nerve and alpha motor neuron activity, leading to muscle contraction and resistance to stretching [21]. In other words, in stimulation with deqi, acupuncture directly stimulates and activates muscle spindles, causing a stretch reflex that causes the muscles to contract.

Involuntary muscle contraction occurs not only by the activation of receptors such as muscle spindles, but also by the action on the nerves connected to the receptors. In a study in which acupuncture needles were applied directly to the median nerve while ultrasound was performed on the forearm, deqi was induced in 85 of 97 patients [22], suggesting that similar responses can be elicited by directly stimulating major nerves. ST36 was defined as being directly above the anterior tibial artery in this study, and the depth at which the subject’s anterior tibial artery was located was 27.7±2.2 mm. The anterior tibialis muscle, which dorsiflexes the ankle joint, is located between the acupuncture needle and the anterior tibial artery. The deep peroneal nerve is parallel to the anterior tibial artery. Hence, determining whether acupuncture causes receptors in the muscle to fire or directly stimulates a major nerve is difficult. The mechanism underlying the increase in myopotential caused by deqi is unclear, but myopotential is a useful objective assessment indicator.

The AWD group showed a higher HR 30 s after stimulation than the AOD group. However, 90 s after acupuncture stimulation, both groups presented a decrease in HR relative to the HR-pre. Therefore, our second hypothesis was not supported, as no increase in HR was observed immediately after deqi occurred. However, the HR immediately after stimulation in the AWD group had different characteristics from those in the AOD group. Temporary excitation of the cardiac sympathetic nerves due to the transient stress response could have inhibited the decreased HR when acupuncture was performed. Numerous studies have indicated that acupuncture reduces the HR [11,13,23–28], caused by somatic autonomic reflex. [11]. This reflex is triggered by the stimulation of nociceptors in the body, which then traverse the C fibers to reach the brainstem and activate the vagus nerve [24]. This study also revealed that in the AOD group, a continuous decrease in the HR occurred immediately after stimulation. This finding supports the results of previous studies. Conversely, in the AWD group, HR did not show a substantial decrease until 60 s after stimulation, with a significant reduction observed at 90 s. When deqi occurred, a transient response in the HR was observed, which was distinct from that observed in the absence of deqi. This seems to be because deqi activates not only somatic autonomic reflexes but also other reflexes and physiological responses.

The first step for elucidating the mechanism of deqi was to find an objective method for detecting it; therefore, we focused our investigation on EMG and ECG findings. The results of this study revealed that when deqi occurs, two characteristic immediate responses are obtained from EMG and ECG: First, the myopotential increases; and second, deqi inhibited the acupuncture-induced decrease in HR. This characteristic response is an important parameter that objectively determines the occurrence of deqi. Furthermore, when evaluated in combination rather than independently, these two characteristics could provide strong evidence for the occurrence of deqi.

### Limitations

Three simulations were performed on 16 subjects during a one-week washout period, yielding data for 48 sessions. The sample sizes differed widely between the two groups, as only six sessions of patients without deqi were obtained. This difference in sample size did not significantly affect the examination of differences between cases with and without deqi. However, to develop an objective index of deqi in the future, increasing and aligning the sample sizes is necessary.

## Conclusion

The EMG hypothesis was that myopotentials would increase during deqi; our results showed that the myopotential significantly increased when deqi occurred, indicating that micro-neural activity was evoked. This finding supports those of previous studies. The hypothesis on HR was that deqi would induce a stress response that would result in a transient increase in HR; our results did not support this hypothesis, as the HR did not increase immediately after deqi onset. However, significant difference in HR was identified immediately after stimulation with deqi compared to stimulation without deqi. Using EMG and ECG, we found that deqi caused an increase in myopotential and inhibited the decrease in HR usually found after acupuncture. These two characteristics enable an objective evaluation of the occurrence of deqi. We investigated whether characteristic responses were obtained from EMG and ECG when deqi occurred, and the results of this study should be used in future research to discriminate the occurrence of deqi.

## Acknowledgments

This work was supported by the JST and the establishment of university fellowships for the creation of science and technology innovation (Grant Number JPMFS2146). We would like to thank Editage (www.editage.jp) for English language editing.

